# Thermal responses of an emerging temperate mosquito reshape arboviral transmission risk

**DOI:** 10.64898/2026.04.14.718355

**Authors:** R Bahrami, D Da Re, A Khorramnejad, DD Gingell, M Brustolin, R Müller, C Damiani, M Bonizzoni, MV Mancini

**Affiliations:** Department of Biology and Biotechnology, University of Pavia, Pavia, Italy; Research and Innovation Centre, Fondazione Edmund Mach, San Michele all’Adige, Trento, Italy; School of Biosciences and Veterinary Medicine, University of Camerino, CIRM Italian Malaria Network, Camerino, Macerata, Italy; Unit of Entomology, Department of Biomedical Sciences, Institute of Tropical Medicine, Antwerp, Belgium

**Keywords:** Invasive species, thermal performance, climate change, arboviruses

## Abstract

Global warming is known to increase arboviral disease risk by enabling the expansion of tropical vectors such as *Aedes aegypti* and *Aedes albopictus*. However, whether climate-driven shifts in seasonal dynamics within temperate regions, such as warmer springs and shorter winters, create thermal settings favourable for additional vectors, which can reshape disease risk maps, is poorly understood. To answer this question, we used the invasive temperate mosquito species *Aedes koreicus* and tested its thermal developmental resilience, along with its vector competence for dengue and chikungunya viruses. We observed significant phenotypic variation in thermal tolerance across life stages, which translates into distinct thermal performance curves for life-history traits, ultimately shaping overall mosquito fitness. We further found that temperature effects are virus-specific, with differential impacts on infection and transmission between dengue and chikungunya viruses. These stage- and pathogen-specific responses generate carry-over effects across life stages, indicating that thermal responses may be constrained by life-history trade-offs rather than be defined by a single thermal optimum.

Our results highlight the need to move beyond static, species-based risk assessments toward mechanistic frameworks that integrate thermal biology across life stages and vector-pathogen systems. As climate change increasingly reshapes seasonal structure rather than simply elevating mean temperatures, cold-adapted and temperate vectors such as *Ae. koreicus*, provide critical model systems to understand how transmission risk emerges outside mosquito classical thermal optima, calling for a broader paradigm shift in how global change and vector-borne disease risk are conceptualised.

## Introduction

The rapid geographic expansion and escalating global transmission of arboviral diseases transmitted by *Aedes* spp. mosquitoes is one of the most dynamic and consequential public health challenges of the past half-century (World Health Organization, 2025). Current approaches to understanding and predicting transmission risk are largely heat-centric, often assuming that warming primarily increases risk by enabling tropical vectors such as *Aedes aegypti* and *Aedes albopictus* to expand into new regions (Craig et al., 1999; Lafferty K.D, 2009; Martens et al., 1997; Parham et al., 2010).

However, this perspective overlooks a critical dimension of climate change in temperate systems: shifts in seasonal structure, such as milder winters and earlier springs, can move species and actively reshape the temporal dynamics of vector activity, survival, and reproduction. Emerging evidence suggests that climate-driven changes in seasonality can also generate novel ecological opportunities in cooler environments, including the expansion of suitable habitats at higher elevations and the extension of activity periods into previously inhospitable seasons (Crowl et al., 2008; Early et al., 2016; Rabitsch et al., 2017; Blaha et al., 2026). These dynamics are likely to disproportionately favour temperate species with the greater tolerance to cold conditions, whose responses cannot be inferred from knowledge generated from tropical vectors, highlighting the limitations of one-size-fits-all assumptions across species.

Within this context, *Aedes koreicus*, an emerged invasive mosquito in Europe, provides a valuable opportunity to address this gap, as its tolerance to colder conditions allows us to investigate processes beyond the typical thermal niche of tropical vectors. Since its first detection in Belgium in 2008, *Ae. koreicus* has rapidly expanded across multiple countries and environmental gradients, including cooler regions and higher altitudes (173–12500 m a.s.l.)(Versteirt et al., 2012; Ballardini et al., 2019; Montarsi et al., 2013). Its ability to overwinter and remain active under relatively low temperatures suggests a thermal niche that extends beyond that of the main European arboviral vector *Ae. albopictu*s (Marini et al., 2019; Da Re et al., 2026).

*Ae. koreicus* is also a key system to investigate the potential epidemiological relevance of cold-adapted mosquitoes because its vector competence to several arboviruses, such as chikungunya (CHIKV) and Zika viruses has been proven (Ciocchetta et al., 2017, 2018; Jansen et al., 2021). However, whether the species’ vector competence is shaped by temperature is poorly understood, with current studies varying in methodological approaches and often relying on simplified assumptions (Ciocchetta et al., 2017, 2018; Jansen et al., 2021). More broadly, current approaches tend to treat thermal responses as uniform across life stages and pathogens (Rebolledo et al., 2021), potentially overlooking critical mismatches that shape both ecological performance and transmission dynamics as thermal conditions experienced during development are known to influence adult performance (Carlassara et al., 2024; Parmesan, 2006;Vanslembrouck et al., 2024)

Using *Ae. koreicus* as a model to examine processes beyond the thermal niche typically considered for tropical vectors, we test how temperature influences *Ae. koreicus* fitness and vector competence for CHIKV and dengue virus (DENV). Based on the thermal performance theory, we hypothesise that (i) thermal performance varies non-linearly across temperature gradients, with distinct optima and limits across life-history traits; (ii) temperature-dependent variation in juvenile development and adult fitness generates effects on relevant traits, including blood-feeding behaviour; and (iii) vector competence for chikungunya and dengue viruses is modulated by temperature in a virus-specific manner.

## Materials and methods

### Establishment of *Ae. koreicus* colony from field-collected individuals

Mosquito larvae were collected in September 2022 from artificial containers in Feltre (Belluno Province, Veneto Region, Italy; 46.0139823 N, 11.8971688 E) using a standard dipping method and transported to the entomological laboratory of the University of Camerino. Larvae (N=600) were reared in the insectary of the University of Camerino at 24 ± 0.5 °C, 70% relative humidity and 14:10 h light: dark photoperiod, using the original field-collected breeding water, and were fed with TetraMin flakes until the L4 stage. Specimens were morphologically and molecularly identified as previously described (Montarsi et al., 2013; Schneider et al., 2016).

Newly emerged adults were further morphologically confirmed (Montarsi et al., 2013) and maintained in BugDorm-6M620 (Insect Rearing cages), with continuous access to 10% sucrose solution. Blood meals were provided 15 days post-emergence, and eggs were collected on white oviposition plates lined with Mosquito Egg Collection Paper EntoWiz (Labitems).

Eggs were maintained at 24 °C under standard laboratory conditions, as previously mentioned, and allowed to quiesce for 5-7 days before hatching (Ciocchetta et al., 2017). Initial attempts to establish the colony using standard hatching protocols for *Aedes* species resulted in low hatching rates (Williges et al., 2008). To optimise hatching success and identify the optimal conditions, eggs were quiesced for 10, 14, 20, 30, and 40 days post-oviposition at at 24 ± 0.5 °C and 70% relative humidity. To induce hatching, three methods were tested for each storage duration: (i) full hypoxia, (ii) partial hypoxia, and (iii) partial hypoxia preceded by an overnight cold exposure at 4 °C. For each treatment combination (quiescence time × hatching method), 100 eggs were used, with two independent biological replicates. In the full hypoxia treatment, eggs were submerged in dechlorinated water and exposed to vacuum pressure for 30 minutes using a polycarbonate vacuum desiccator (Brand™, Wertheim, Germany). In the partial hypoxia treatment, eggs were floated in sealed containers containing dechlorinated water, leaving approximately 50% of the total volume as air space to create reduced oxygen conditions without vacuum application. For the cold-exposure treatment, eggs were first maintained at 4 °C overnight and subsequently subjected to the partial hypoxia hatching protocol.

After treatment, eggs were kept submerged in dechlorinated water for 24 hours, after which the number of first-instar larvae was recorded. Unhatched eggs were then dried and re-submerged for two additional hatching cycles following the same procedure. Five generations (G5) after establishment, eggs of the *Ae. koreicus* colony were shipped to the University of Pavia, where egg quiescence was standardised to 20 days (Figure S 1A), before rearing at standard conditions, as previously described. Eggs are hatched using the partial hypoxia method described above. Larvae are reared in plastic containers (19 × 19 × 6 cm) at a controlled density of approximately 200 larvae per litre of dechlorinated water to reduce intraspecific competition. Larvae are fed daily fish food (TetraMin, Tetra, Germany). Adults are maintained in mesh cages (BugDorm; 60 × 60 × 120 cm) and provided with a 10% sucrose solution via cotton pads *ad libitum*. Female mosquitoes are offered human blood (Centro Trasfusionale, IRCC Policlinico San Matteo, Pavia, Italy) using a membrane feeding system (Hemotek, UK). Blood meals are offered 11 days post-emergence. To determine the duration of the gonotrophic cycle and optimise oviposition timing for colony maintenance, we monitored female fecundity following two consecutive blood meals. Non-engorged females were removed after blood feeding, whereas fully engorged females were retained and allowed to oviposit on damp filter paper starting at three days post-feeding. Eggs were collected and counted daily to quantify fecundity and determine the temporal pattern of oviposition. If no eggs were laid for two consecutive days, a second blood meal was offered to the same cohort, and egg deposition was monitored daily. This procedure was performed in two independent biological replicates and served to characterise the gonotrophic cycle of *Ae. koreicus* and to define the optimal oviposition timing at 5 days post blood meal (DPB) for routine colony maintenance (Figure S 2A).

### Measurements of life history traits

We reared *Ae. koreicus* under six constant temperatures (16, 20, 24, 28, 32, and 36°C), which reflect the range of average environmental conditions at the original collection site in Feltre (Belluno Province, Veneto Region, Italy) of the mosquito population (ARPAV, 2023). We kept relative humidity (70 ± 5%) and photoperiod (14:10 h light: dark) constant in climate-controlled chambers to isolate the effects of temperature. Fitness experiments were performed on cohorts of mosquitoes of G7 from colony establishment.

For each temperature treatment, we processed batches of 150–200 eggs across four biological replicates using the optimised hatching protocol described above. Hatching rates were assessed 24 hours after submersion by counting the number of first instar larvae. Unhatched eggs were subjected to the floating hatching protocol described previously, involving two additional hatching rounds to maximise larval recovery. Hatching rates represent the proportion of the total number of emerged larvae across all three hatching rounds relative to the total number of eggs. Batches of hatched larvae from each round were transferred to plastic trays containing 200 mL of dechlorinated water and fed one-quarter of a Tetramin™ tropical fish food tablet daily. Larvae were reared under their assigned temperature regime and monitored daily until pupation to measure larval developmental time (LDT). Pupae and adults were counted to evaluate pupation rate (number of pupae over total number of larvae), pupal developmental time (PDT), and adult emergence rate (number of adults over total number of pupae). We also calculated larval-to-adult development time (hereafter called developmental speed) and larval-to-adult survival (hereafter called developmental success)(Dennington et al., 2024). Developmental speed was calculated as the inverse of the median time from first-instar larva to adult emergence (day^− 1^). Developmental success was analysed as the number of adults emerging from a known number of introduced larvae for each replicate. Upon eclosion, adults were sexed and counted to determine the sex ratio. For adult longevity, 10 males and 10 females in four biological replicates per thermal regime were placed into BugDorm cages (12 × 12 × 12 cm) and maintained under their respective temperature regimes. Mortality was recorded daily until 80% of mosquitoes had died. Only for the adults reared at 16°C, longevity was monitored until the last individual died, allowing proper handling of censored observations. We randomly sampled a total of 30 females per treatment to measure their wing length, which was used as a proxy for adult size (Robert et al., 1998). The right wing was dissected and measured from the alular notch to the apical margin, excluding fringe scales, using an Olympus CKX53 inverted microscope equipped with CellSens Standard software. Females were offered a blood meal 10 days post-emergence to estimate their reproductive capacity. Fully engorged females were isolated and maintained individually for a fixed period of 6 days post-blood meal (DPB), after which ovaries were dissected and the number of mature oocytes (defined as egg load) per female was counted using an Olympus SZX10 stereo microscope.

### Cold Stress Assay

We measured cold-induced knockdown time, defined as the time required for mosquitoes to become immobilized following exposure to 4 °C. It was assessed in 11-day-old mosquitoes reared at either 24 °C or 28 °C and maintained on a sugar diet to eliminate potential confounding effects related to blood-digest ion-induced stress. Adult males and females were briefly chilled and placed individually into 2 mL microcentrifuge tubes and left for an additional 4 hours at their respective rearing temperatures before being transferred to a 4 °C climate chamber, and their movements were monitored every 5 minutes until complete knockdown. We tested groups of 15 sugar-fed mosquitoes of the same sex and rearing condition, for a total of 30 mosquitoes per sex and per thermal treatment.

### Measurement of blood feeding behaviour and blood digestion

We monitored female blood-feeding behaviour under either constant 24 °C or 28 °C. For each thermal regime, mosquitoes were reared in colony cages (60 × 60 × 120 cm) at a 3:1 male-to-female ratio to ensure mating. A preliminary blood meal was offered at 0–3 days post-emergence (DPE) while mosquitoes were still maintained in the larger colony cages; however, no females fed at this stage. Individuals were therefore maintained in these cages to allow mating prior to subsequent feeding attempts. Five-seven days post-emergence, 70 females were transferred to smaller cages (45 × 45 × 45 cm) and offered blood for 90 minutes under their respective rearing temperatures to avoid behavioural changes due to thermal mismatch. Subsequent blood meals were offered at 9–11, 15–17, and 20–22 DPE. After each feeding event, engorged females were recorded and removed from the cages. After a blood meal to 9-11-day-old females, 50 engorged females were sampled immediately to quantify blood intake, measuring haemoglobin using Drabkin’s reagent (Sigma-Aldrich, Cat. D5941). One vial of Drabkin’s reagent was dissolved in 1 L of distilled water, and 0.5 mL of 30% Brij-35 (Sigma-Aldrich, Cat. B4184) was added to obtain a 1× working solution, following the manufacturer’s instructions. For each sample, 500 μL of Drabkin’s reagent was added to the dissected abdomen and homogenised for approximately 30s using a polypropylene pellet pestle (KIMBLE, Cat. 749521-0500). Samples were incubated at room temperature, centrifuged at 500 × g for 1 min, and the supernatant was transferred to a new tube. To quantify haemoglobin, 200 μL of each mosquito sample and a serial dilution of the same blood used for feeding (0, 0.2, 0.4, 0.6, 0.8, and 1.0 μg μL^− 1^) were loaded into a 96-well borosilicate plate. Absorbance was measured at 540 nm. Haemoglobin concentration was used as a proxy for blood intake per individual mosquito.

For each temperature, females were sampled at 12, 24, 48, 72, and 96 h post-blood meal to quantify trypsin-like activity. At each time point, 30 females were analysed, consisting of three biological replicates of 10 females each. Females were briefly chilled at 4 °C, and their abdomens dissected on ice to assess trypsin-like activity following the protocol described (Gulia-Nuss et al., 2012). Briefly, each midgut was homogenised in 100 μL of extraction buffer (20 mM Tris, 20 mM CaCl_2_; Sigma-Aldrich; pH 8.0) using a disposable pestle. Homogenates were centrifuged at 14,000 × g for 2 min at 4 °C. Supernatants were collected and stored at −80 °C until analysis. For each sample, 5 μL of supernatant was added to 100 μL of 4 mM Nα-benzoyl-L-arginine-p-nitroanilide (BApNA; Sigma-Aldrich) in a 96-well plate. Plates were incubated at 37 °C for 10 min, and absorbance was measured at 405 nm using a CLARIOstar microplate reader (BMG Labtech). Trypsin-like activity was quantified using a standard curve generated with bovine serum trypsin (Sigma-Aldrich; 20 μg) and BApNA standards (8.96, 4.48, 2.24, 1.12, 0.56, 0.28, and 0.14 mM).

### Measurement of mosquito vector competence to DENV and CHIKV

We tested *Ae. koreicus* vector competence to DENV and CHIKV using females reared either at 24 °C and 28 °C. For each thermal treatment, batches of 80 females aged 9–11 days were sorted and starved for 48 h prior to exposure to an artificial infectious blood meal. Experimental groups were fed on virusspiked (final titer of 1*10^7 focus-forming units per mL (FFU/mL) for either CHIKV or DENV-2) human blood (Belgian RedCross) via the Hemotek feeding system (SP6W1-3; Hemotek Ltd, Blackburn, UK) using a collagen membrane (MEM5; Hemotek Ltd) at 38 °C for 45 min. We used CHIKV strain LR2006 and DENV-2 strain 15091930. Mosquitoes were assigned to four experimental groups based on temperature and virus exposure: 24 °C–CHIKV(n=60), 24 °C–DENV(n=79), 28 °C–CHIKV(n=75), and 28 °C–DENV(n=63) (Table S1). After infection, mosquitoes were sampled 7-, 14-, and 21-days post-infection (DPI) to assess infection rate (IR), disseminated infection rate (DIR), and transmission efficiency (TE). Briefly, infection rate (IR) was defined as the proportion of virus-positive mosquito bodies among the total number of blood-fed females. Disseminated infection rate (DIR) was defined as the proportion of mosquitoes with infected legs among those with infected bodies. Transmission efficiency (TE) was defined as the proportion of mosquitoes with infectious virus detected in saliva among the total number of blood-fed females. A salivation assay was performed as previously described (Heitmann et al., 2018, Brustolin et al., 2024). Briefly, mosquitoes were induced to salivate for 20 min. After salivation, bodies and legs were collected separately. Samples were homogenised in mosquito diluent consisting of phosphate-buffered saline (PBS 1×) supplemented with 20% fetal bovine serum (FBS), 50 μg mL^− 1^ penicillin–streptomycin, 50 μg mL^− 1^ gentamicin, and 2.5 μg mL^− 1^ amphotericin B. The presence of infectious virus particles in bodies, legs, and saliva was assessed using a focus-forming assay (FFA) (Brustolin et al., 2023). For all collected samples, 30 μL of each saliva or tissue homogenate was inoculated onto confluent cell monolayers. Dengue virus samples were incubated on Vero cells, whereas chikungunya virus samples were incubated on BHK cells. Following infection, virus detection was performed using virus-specific primary antibodies, including the monoclonal anti-CHIKV E2 envelope glycoprotein clone CHK-48 (BEI Resources, Manassas, VA, USA) for CHIIKV, and the anti-DENV Virus Complex Antibody, clone D3-2H2-9-21 (Chemicon, Merck KGaA, Darmstadt, Germany) for DENV.

### Statistical analyses of fitness data and viral titers

The Shapiro–Wilk test was used to assess the normality of data distributions (Ghasemi & Zahediasl, 2012). Depending on the outcome, parametric analyses of variance (ANOVA) or non-parametric tests (Kolmogorov–Smirnov, Mann–Whitney, or Kruskal–Wallis) were applied accordingly (Ramachandran & Tsokos, 2020). Statistical significance was set at p ≤ 0.05. Differences across temperatures in egg hatching rate, pupation rate, adult emergence rate, LDT, PDT, and adult sex ratio were analysed using two-way ANOVA followed by Tukey’s multiple comparison test (Lee & Lee, 2018). Larval and pupal development rates (LDR and PDR) were calculated as the multiplicative inverse of LDT (1/LDT) and PDT (1/PDT), respectively. Their ratio (LDR/PDR) was analysed using regression models across developmental temperatures (Folguera et al., 2010). Comparisons of mosquito longevity were carried out with the Kaplan–Meier survival analysis and log-rank (Mantel–Cox) test (Kishore et al., 2010). These analyses were performed using Prism 8 (GraphPad Software). Since a proportion of individuals were still alive at the end of the experiments, they were right-censored. To verify that the censoring protocol did not introduce systematic bias in longevity estimates, we used the 16°C dataset as a validation case, as it was the only temperature at which all individuals were observed to death with no censoring applied. We artificially applied the experimental censoring rule to these fully observed data by identifying the day on which 80% of deaths had occurred within each sex (cutoff day: females = 74 days, n = 40 deaths out of 50; males = 51 days, n = 29 deaths out of 36) and right-censoring all surviving individuals beyond that day. We then compared mean longevity between the fully observed and artificially censored datasets using a linear mixed model (Gaussian family, identity link) with censoring status (full vs. censored) as a fixed effect and replicate as a random intercept, fitted with the *glmmTMB* package (Brooks et al., 2017). Model fit was assessed using DHARMa scaled residual diagnostics (Hartig et al., 2022).

Viral outcomes were analysed using complementary approaches depending on the response variable. IR, DIR and TE were calculated as observed proportions with a Chi-square test. CHIKV infection and dissemination titers were analysed using hurdle generalized linear models (GLMs), composed of a zero-truncated negative binomial GLM fitted using the R package *glmmTMB* (Brooks et al., 2017) to account for overdispersed count data with excess zeros. Zero counts were treated as true uninfected individuals with the model separating infection probability (zero component) from infection intensity among infected individuals (conditional component). Temperature and days post-infection (DPI), and their interaction, were included as fixed effects. Model fit was assessed using Akaike Information Criterion corrected for small sample sizes (AIC) and DHARMa (Hartig et al., 2022). Estimated marginal means and pairwise comparisons were obtained using the *emmeans* package with Holm correction for multiple testing. Infection probabilities were calculated directly from observed data. In contrast, CHIKV transmission data sets and DENV datasets (infection, dissemination and transmission) contained a high proportion of zero values and highly skewed distributions, which prevented reliable model fitting. Therefore, DENV data were analysed using non-parametric methods (Kruskal–Wallis tests with pairwise Wilcoxon comparisons).

### Design of thermal performance curves

Thermal performance curves (TPCs) were fitted separately for three life-history traits of *Ae. koreicus*: developmental speed, developmental success, and adult longevity (male and female) using R v4.5 (R Core Team, 2024). A frequentist framework was adopted, and model structures were selected according to the statistical properties of each response variable. We selected model families appropriate to the error structure of each trait, rather than applying a uniform framework across all traits. Model selection was based on the AIC (Burnham & Anderson, 2002), and residual diagnostics were evaluated using the *DHARMa* package (Hartig, 2022), which generates simulation-based scaled residuals from the fitted model distribution.

Thermal performance for developmental speed was described using a Brière-2 function fitted by nonlinear least squares via *nls*.*multstart* package (Briere et al., 1999; Reuss et al., 2018; Padfield et al., 2020). Candidate functions were compared using AICc (Yin et al., 1995). Models were implemented using *rTPC* package (Padfield et al., 2021). The lower critical thermal limit (CT^min^) was taken directly from the estimated *t*^min^ parameter of the Brière-2 function, which represents the temperature at which the development rate mathematically returns to zero. The upper thermal limit (CT^max^) was set to 36°C based on empirical observation — no development was recorded at or above this temperature — as the Brière *t*^max^ parameter was boundary-constrained at 36°C due to the limited temperature range of the experiment. The thermal optimum (T^opt^) was defined as the temperature at which the fitted curve reaches its maximum, extracted from a fine-resolution prediction grid (0.2°C intervals). Confidence intervals were generated by residual bootstrap resampling (999 iterations).

Developmental success was modelled as a binomial outcome (number of individuals that reached adulthood vs number of individuals that failed to eclose) in a GLMM model with quadratic temperature as a fixed effect and replicates’ ID as a random effect. Models were fitted using the *glmmTMB* package (Brooks et al., 2017) and the thermal optimum was derived analytically from the quadratic coefficients as Topt = −β1 / (2β2), where β1 and β2 are the linear and quadratic temperature coefficients, respectively. Predictions and 95% confidence intervals were computed on the logit scale and back-transformed to the response scale to ensure bounds remained within [0, 1]. Model selection was based on the AIC and likelihood ratio test comparisons, while residual diagnostics were evaluated using the *DHARMa* package (Hartig, 2022).

Adult longevity data consisted of individual-level time-to-event records for female and male mosquitoes across five experimental temperatures. The adults’ true longevity exceeded the recorded observation day, and using mean longevity per replicate as a response variable in nonlinear least squares regression would have been inappropriate as it underestimates true longevity in the presence of censoring (Clark, 2003). We therefore used a parametric survival modelling approach that correctly incorporates censored observations through the likelihood function. Parametric accelerated failure time (AFT) models were fitted using the *flexsurv* package (Jackson, 2016), with temperature entered as a quadratic fixed effect to capture the unimodal thermal response. Male and female longevity were analysed as separate models, as thermal sensitivity may differ between sexes, and three candidate distributions were compared (Weibull, gamma, lognormal) by AIC. Mean expected longevity at each temperature was derived from the fitted model using the *summary*.*flexsurvreg* function with type = “mean”, with 95% confidence intervals computed via the delta method.

## Results

### *Ae. koreicus* can develop between 13.2 and 36°C

We measured *Ae. koreicus* life-history traits across a gradient of temperatures from 16 to 36 °C (Figure 1A). We saw pupation rate (p=0.03, F _4-11_= 3.72, ANOVA) and juvenile developmental time both LDT (p<0.0001, F_4-12_= 95.87, ANOVA) and PDT (p<0.0001, F_4-12_= 30.94, ANOVA), reducing with increasing temperatures, with a drastic change at 32°C (Figure 1B, F, and G). In contrast, hatching and adult emergence rates did not differ significantly across temperatures, except at 36 °C, where larval emergence was extremely limited and inconsistent across replicates. Only two larvae were observed in a single replicate, and both died within 24 hours, indicating that this temperature exceeds the upper thermal limit for successful early larval development under our experimental conditions.

**Figure 1.**
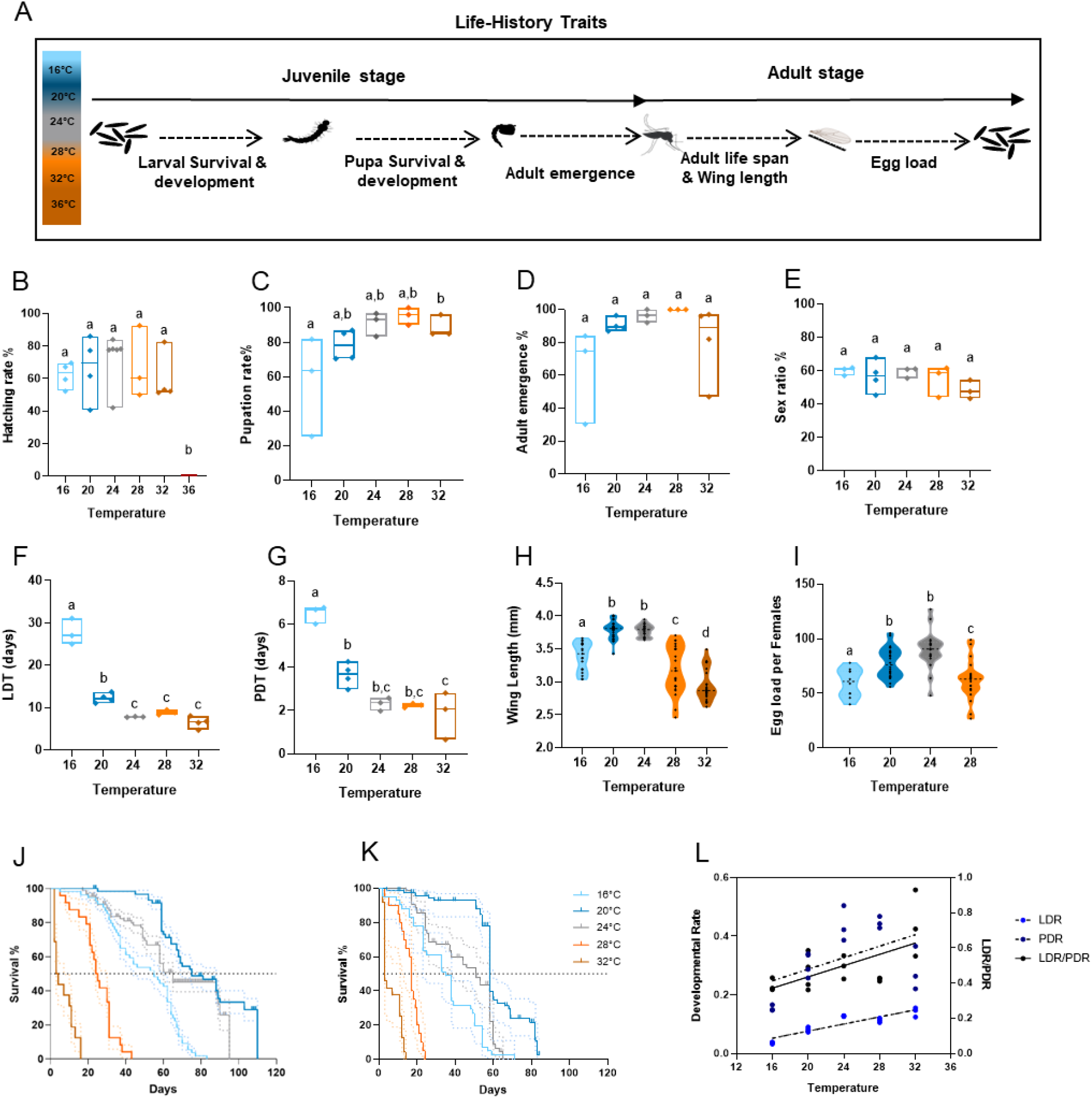
Temperature effects on life-history and morphological traits of *Ae. koreicus* during juvenile and adult stages. (A) Experimental design and life-history tracking across constant temperature treatments, from egg hatching through larval and pupal development to adult emergence and reproduction, were used to quantify temperature-dependent traits. (B) Hatching rate, including the 36 °C treatment, at which only two larvae emerged; this temperature was excluded from subsequent fitness traits. (C) Pupation rate. (D) Adult emergence rate. (E) Sex ratio of emerged adults. (F) Larval developmental time (days). (G) Pupal developmental time (days). (H) Female wing length as a proxy for adult body size. (I) Egg load in the ovaries of adult females; data at 32 °C are not shown due to reduced adult lifespan. (J) Female survival and (K) Male survival. (L) The isomorphy rule is expressed as the ratio of larval to pupal developmental time (LDR/PDR) across temperatures. Boxplots and violin plots represent the distribution of observed values, with individual points showing replicate measurements. Different lowercase letters indicate significant differences among temperature treatments (post hoc tests, p < 0.05).

Given the observed differences in developmental speed, we tested the hypothesis of developmental isomorphy, which is considered a general rule for ectotherms (Jarošík et al., 2004). Regression analysis revealed a positive relationship between LDR and developmental temperature. A similar trend was observed for PDR. However, LDR and PDR slopes were significantly different, with a significant positive regression with respect to temperature (*b* = 0.003299) (Figure 1L). The increase in the LDR/PDR ratio indicates that *Ae. koreicus* larval development is more constrained by temperature than pupal development.

In the TPC analyses for developmental success, initial binomial models exhibited substantial overdispersion (Pearson dispersion ratio = 5.5), which was effectively accounted for using a beta-binomial model. Inclusion of a replicate-level random effect did not improve model fit (ΔAICc = 2; LRT p = 1), indicating that variability was adequately captured by the dispersion parameter. We observed that developmental success peaked (Topt at 26.3±0.02 °C). Developmental success declined steeply toward both thermal extremes with an estimated critical thermal minimum (CTmin) of 13.2 ± 2.3 °C (Figure 2A) and observed CTmax at 36°C.

**Figure 2.**
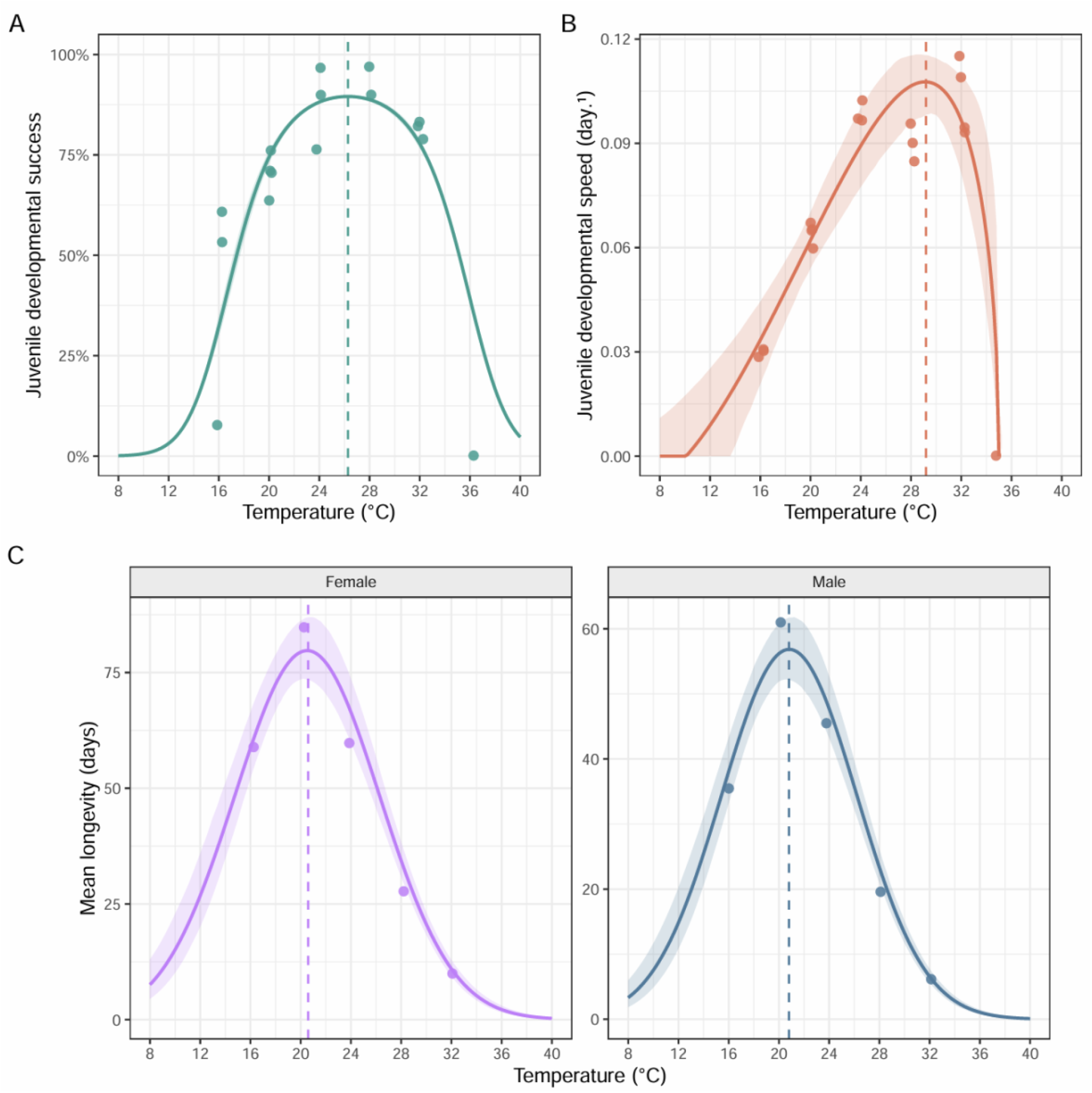
Thermal performance of life-history traits in *Ae. koreicus*. (A) Developmental success (probability of larval survival to the adult; Topt =26.8°C). (B) Developmental speed (Topt = 29.6 °C). (C) Probability of female survival (Topt = 20.6 °C). (D) Probability of male survival (Topt = 20.8 °C). Points represent observed values, solid lines show fitted thermal performance curves, and shaded areas indicate 95% confidence intervals where available. Vertical dashed lines denote estimated thermal optima.

For developmental speed, the Brière-2 model provided the best fit among candidate models (ΔAICc > 13), capturing the unimodal relationship between temperature and development rate. Developmental speed increased with temperature up to a maximum of 0.101 day^− 1^ at 29.6±0.95 °C before declining toward the upper thermal limit (Figure 2B; Table S2).

### Adults have a thermal optimum of 20°C, where longevity extends past two months

Adult sex ratio remained stable across the tested temperatures, ranging from 48.4% in 32°C to 59% in mosquitoes reared at 16°C, indicating no detectable thermal effect in sex allocation (Figure 1E). In contrast, wing length showed a clear temperature-dependent pattern. The largest females emerged at 20°C, whereas both cooler (16 °C) and warmer (32 °C) conditions produced significantly smaller individuals (Figure 1H; p < 0.0001 F_4-117_=72.984, ANOVA) a different thermal profile. Egg load peaked at 24 °C and declined toward both thermal extremes in our experimental set up (p<0.0001 F_3-73_=13.66, ANOVA) (Figure 1I). Adult survival declined progressively with increasing temperature, with median lifespan highest at 20 °C (females: 75 days; males: 58 days) and sharply reduced at 32 °C (females: 3.5 days; males: 3 days) (Figure 1J–K). At 32°C, the limited number of surviving females, combined with reduced feeding success, precluded a detailed assessment of ovarian maturation stages and reliable estimates of egg load. This pattern was supported by TPC analyses, through fitting the Weibull distribution for both sexes (ΔAICc > 6.5 over lognormal; gamma failed to converge). The Weibull shape parameter exceeded 2 for both sexes (female: 2.33; male: 2.40), consistent with an increasing hazard rate and actuarial senescence, which estimated an optimal temperature for longevity at 20.6 °C, with predicted mean lifespans of 79.7 days for females and 60.3 days for males (Figure 2 C, Table S2).

### *Ae. koreicus* females prefer to feed on blood at 24°C rather than 28°C

Based on the observed peak of egg production at 24°C, we decided to extend the analysis to other traits that could impact *Ae. koreicus* physiology and vector competence at this temperature, comparing it to 28°C, which represents the standard rearing temperature for tropical mosquitoes (Figure 3A). We saw the peak of blood-feeding activity shifting 2-4 days in relation to the rearing temperature. The peak in blood feeding was observed in 9–11-day old females at 24°C and in 5–7-day-old females at 28°C (Figure 3B). Notably, at their respective peak, the number of blood-feeding females at 24°C was approximately twice than that observed at 28 °C. Haemoglobin quantification showed that females reared at 24 °C intake significantly larger blood meals than those reared at 28 °C (Figure 3C; p = 0.0041, Mann Whitney), resulting in two-fold higher trypsin activity in females reared at 24 than at 28°C (Figure 3D). Accordingly, the time to complete blood digestion was 24h longer for females reared at 24 than those kept at 28 °C (Figure 3E and F). We further examined the cold-induced knockdown time of adults exposed to 4°C after having been reared at 24 °C or 28 °C. For both sexes, we observed that acclimation at lower temperature increased cold resilience. Knockdown time was longer for adults reared at 24°C (45.55 min in females and 36.92 min in males,) than 28°C (33.06 min for females). Females reared at the higher temperature (28 °C) exhibited a shorter knockdown time (33.06 min) compared to those reared at 24 °C (45.55 min) (p<0.0001, Mann Whitney). A similar pattern was observed in males, with individuals reared at 28 °C showing faster knockdown (30.7 min for males) than those reared at 24 °C (36.92 min) (p=0.015, Mann-Whitney) (Figure S3 A-B).

**Figure 3.**
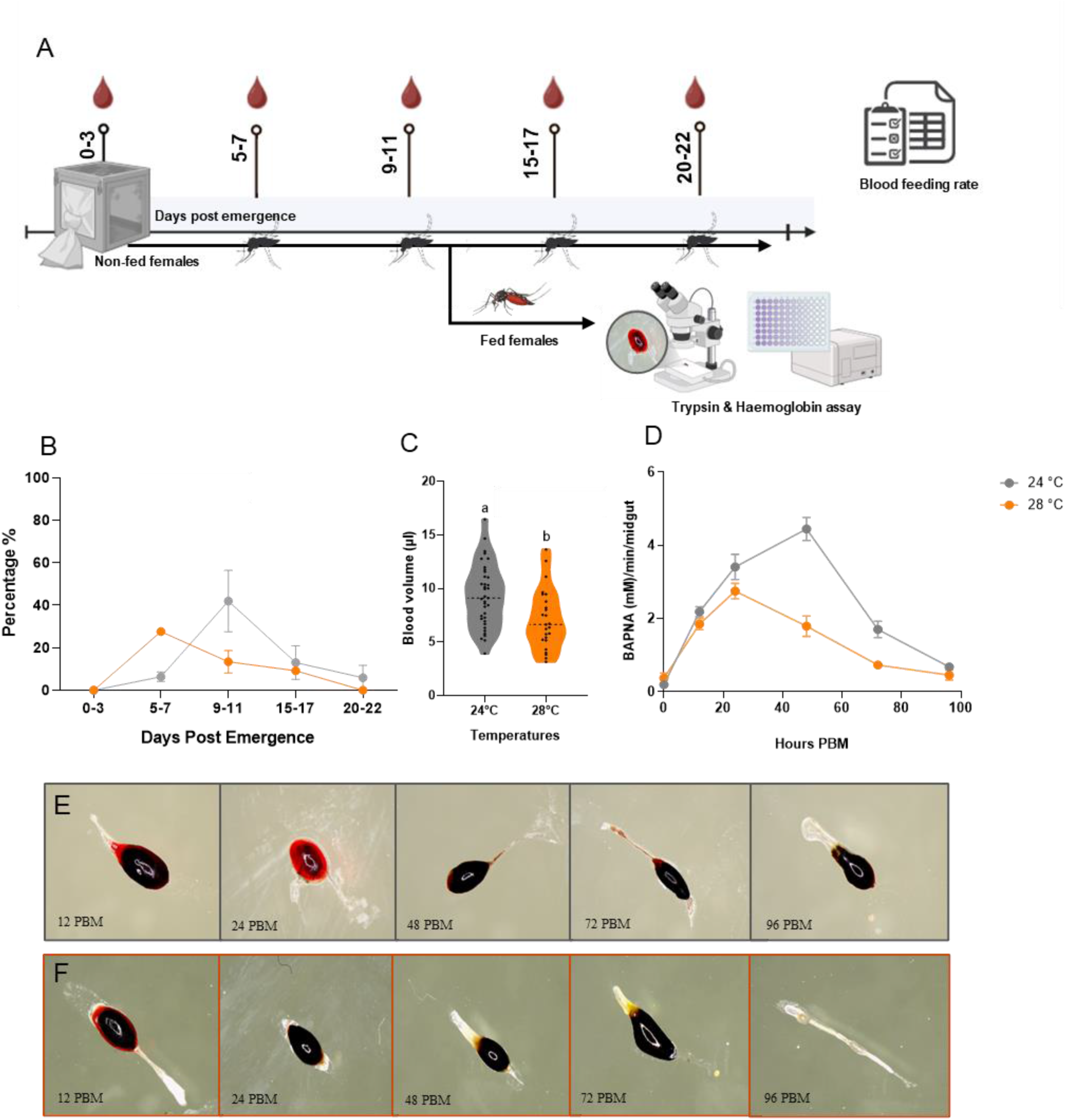
Temperature-dependent blood-feeding traits of *Ae. koreicus* females. (A) Experimental design for females reared at constant temperatures of 24 °C and 28 °C. (B) Comparison of blood-feeding rates of females reared at 24 °C and 28 °C across different time points post-emergence. (C) Comparison of blood meal volume intake by females reared at 24 °C and 28 °C, quantified using a hemoglobin assay. (D) Comparison of blood digestion dynamics under the two thermal regimes, assessed using a colorimetric assay and trypsin activity measurements. (E, F) Representative images of mosquito midguts at different time points post blood meal for females reared at 24 °C (E) and 28 °C (F).

### Vector competence in *Ae. koreicus* is virus-specific and temperature-dependent

We studied the vector competence for both CHIKV and DENV in mosquitoes reared at either 24°C or 28°C by examining viral load 7, 14, and 21 days post-infection (DPI) in whole bodies, legs, and saliva.

In mosquito bodies, CHIKV IR resulted to be temperature- and DPI-independent, reaching above 80% of infected mosquitoes. In contrast, viral titres were influenced by temperature in a time-dependent manner, with a significant temperature × DPI interaction (p=0.045). At 7 DPI, titres were higher at 24 °C than at 28 °C (∼4.7-fold, p=0.0013). This difference was no longer observed at 14 DPI (p=0.739), but re-emerged at 21 DPI, when titres were again higher at 24 °C (∼8.8-fold, p < 0.0001) (Figure 4A). Over time, titres declined at both temperatures, but with different patterns. At 24 °C, the decrease occurred early, between 7 and 14 DPI (∼8-fold, p=0.0001), and then stabilised. In contrast, at 28 °C, the decline was delayed and became more pronounced by 21 DPI (∼7-fold from 7 DPI, p < 0.0001) (Figure 4A). In legs, DIR significantly increased over time only at 24°C (p=0.002,df=2, Chi-square). CHIKV titres were consistently lower at 28 °C across all time points. Differences between temperatures were already evident at 7 DPI (∼4.2-fold, p=0.0137) and persisted through 14 DPI (∼3.9-fold, p=0.0271), increasing further by 21 DPI (>12-fold, p < 0.0001). No significant temperature × DPI interaction was detected. However, titres declined over time at both temperatures, with the strongest reductions observed at 21 DPI(Figure 4B). We detected virus in saliva only 7 DPI in 22% of mosquitoes reared at 24°C, and this percentage decreased to 5% in mosquitoes reared at or 28 °C (Figure 4C).

**Figure 4.**
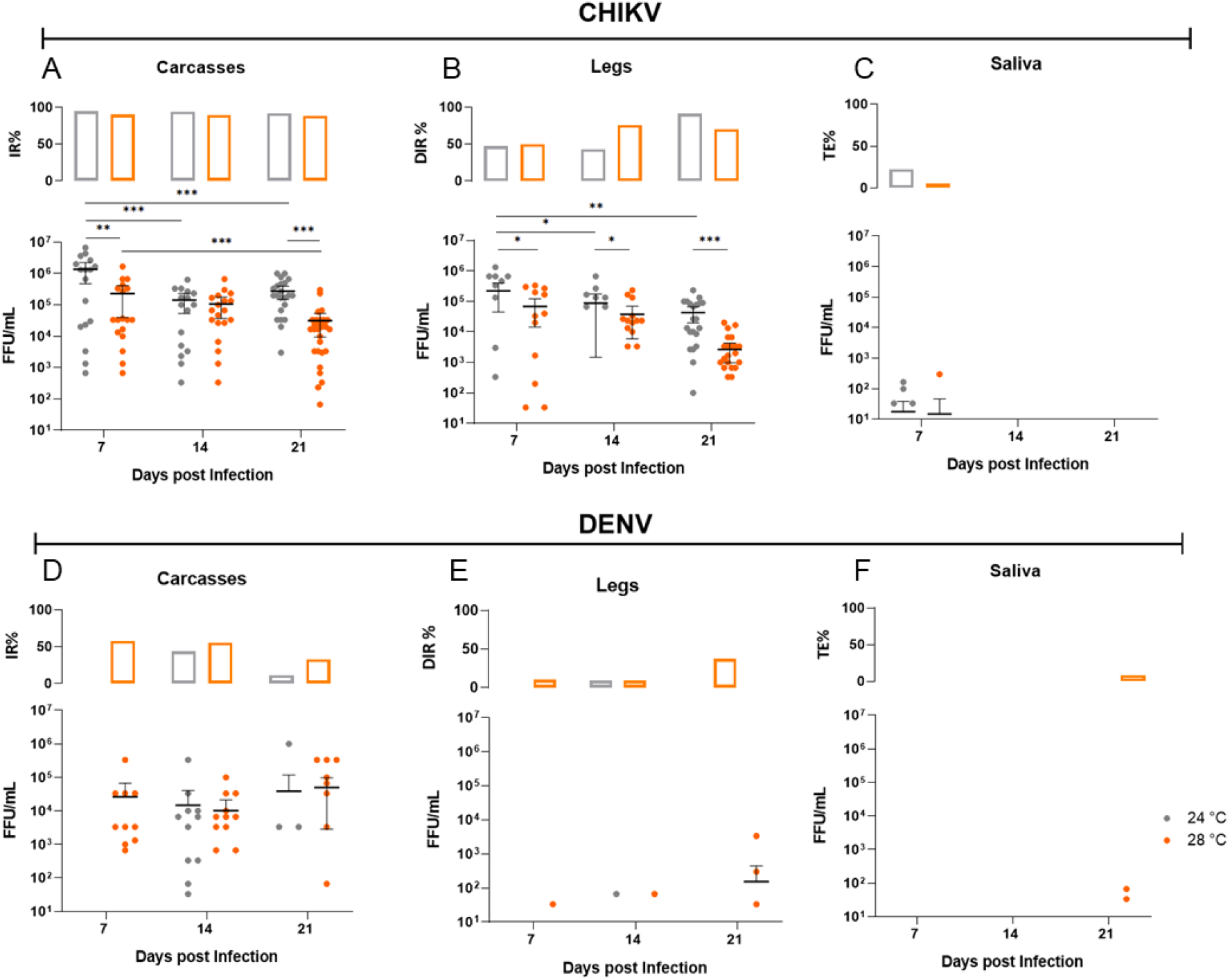
The vector competence of *Ae. koreicus* for CHIKV and DENV under two different thermal regimes. Vector competence of *Ae. koreicus* for chikungunya virus (CHIKV; panels A– C) and dengue virus (DENV; panels D–F). (A and D) Infection dynamics in mosquito carcasses. (B and E) Dissemination dynamics in legs. (C and F) Transmission dynamics in saliva. For each tissue, the upper subpanels indicate the proportion (%) of infected, disseminated, or transmitting mosquitoes, while the lower subpanels show individual viral titers expressed as FFU/mL on a log_10_ scale. Each dot represents a single mosquito; horizontal bars indicate mean ± SEM; IR, infection rate; DIR, disseminated infection rate; TE, transmission efficiency; FFU, focus-forming units;

DENV infection was detected earlier at 28 °C with 50% (IR) at 7 DPI, whereas at 24 °C it was first observed at 14 DPI (40.7%) (Figure 4D). In contrast to CHIKV, DENV viral load remained stable over time and did not significantly differ between temperatures. Dissemination was limited at early time points: at 24 °C, 9.1% of mosquitoes showed positive legs at 14 DPI. At 28 °C, dissemination remained low at earlier time points (10% at 7 DPI; 9% at 14 DPI), but showed an increasing trend reaching 37% at 21 DPI (Figure 4E). Accordingly, DENV was detected in saliva only at the latest time point, 21 DPI, in 8.3% of mosquitoes (Figure 4F).

## Discussion

As climate regimes shift, environmental conditions are being rapidly reshaped, altering species distributions and ecosystem dynamics. While much attention has focused on the expansion of warm-adapted mosquitoes, cooler regions may increasingly be colonised by cold-tolerant invaders. Understanding how such species respond to temperature is therefore critical for predicting their persistence and estimating the risk of arboviral transmission. We addressed these questions using *Ae. koreicus*, a species recently invading continental Europe from temperate Asia. Our results reveal pronounced stage-specific thermal responses in *Ae. koreicus*. Juvenile traits performed best between 13-35°C, a range that overlaps with temperatures commonly experienced during spring and summer in temperate regions (WeatherSpark, 2026). Like other *Aedes* species, developmental success and developmental speed showed different thermal optima (26.8 °C and 29.6 °C, respectively), with developmental speed often having a higher optimum compared to success (Dennington et al., 2024). In contrast, adult fitness traits showed a different thermal profile: longevity collapsed at 32 °C, and this pattern was consistent across both sexes, suggesting proximity to the upper thermal boundary of this species. For females this also represents the threshold for successful completion of the gonotrophic cycle. By applying a thermal performance framework, our results advance previous findings by providing a mechanistic prediction of adult performance, indicating an optimum around 20 °C (Marini et al., 2019; Montarsi et al., 2013). The mismatch between juvenile and adult thermal optima is particularly relevant because the environmental conditions experienced during juvenile development and adult life stage can influence population growth and dispersion, although their relative importance varies across systems (Mordecai et al., 2019; Pawar et al., 2024). Cooler thermal optima for adults compared to juvenile stages likely help explain the distribution of *Ae. koreicus* across the temperate regions of Europe, including its rapid spread in pre-alpine northern Italy (Knight, 1947; Capelli et al., 2011; Marini et al., 2019; Miyagi,1971). In these environments, moderate temperatures can simultaneously support rapid juvenile development and extended adult survival. Importantly, our experiments were conducted on an established field population from northern Italy at the edge of the current distribution range: defining the thermal performance of this population is particularly valuable, as it underscores its potential for further expansion. Whether expanding populations will undergo local adaptation or shift their thermal limits remains unknown.

The thermal profile of *Ae. koreicus* contrasts with that of more extensively studied invasive species, such as *Ae. aegypti* and *Ae. albopictus*, whose performance and spread are often associated with warmer climatic optima (Carlassara et al., 2024). From a comparative perspective, *Ae. koreicus* therefore represents a functionally distinct invader, potentially occupying a cooler segment of the thermal niche space among invasive *Aedes*. Nevertheless, although reduced, the median longevity at 28 °C was comparable to values reported for *Ae. albopictus* (Carlassara et al., 2024; Montarsi et al., 2013b), indicating partial thermal overlap between the species. While studies have reported weak larval competition between *Ae. albopictus* and *Ae. koreicus* (Baldacchino et al., 2017), this overlap in adult thermal performance suggests that competition could occur at later life stages. Consequently, *Ae. koreicus* may persist during cooler periods while overlapping with *Ae. albopictus* during warmer conditions, influencing local population dynamics, seasonal abundance, and temporal niche partitioning in temperate regions. Temperature simultaneously constrains mosquito fitness, physiology, and capacity to support viral replication, making it a key determinant of transmission dynamics (Couper et al., 2024; Mordecai et al., 2019b). In *Ae. koreicus*, we observed clear temperature-dependent differences in behavioural and infection-related traits under ecologically relevant conditions (24 °C and 28 °C). Our results indicate that warmer conditions (28 °C) accelerated blood-meal processing and shifted peak feeding activity, whereas cooler conditions (24 °C) favoured higher feeding success. Reduced feeding at 28 °C may indicate mild thermal stress impacting host-seeking behaviour, a phenomenon observed in other arthropods (Slayi & Jaja, 2025). Higher feeding success at 24 °C, representative of early autumn and late spring conditions in temperate regions, suggests that *Ae. koreicus* could initiate virus transmission early in the season. Our results indicate that *Ae. koreicus* is capable of transmitting both CHIKV and DENV under laboratory conditions, with transmission strongly modulated by temperature. CHIKV transmission occurred at 24 °C, whereas DENV transmission was restricted to 28 °C, indicating distinct thermal thresholds for arboviral transmission in this species. Differences in transmission dynamics between CHIKV and DENV are reflected in infection and dissemination patterns: for CHIKV, relatively high infection and dissemination rates coupled with low transmission efficiency suggest the presence of internal barriers limiting viral release into the saliva, whereas for DENV, low dissemination rates point to a potential midgut escape barrier restricting viral movement to secondary tissues. The earlier detection of CHIKV compared to DENV is consistent with its shorter extrinsic incubation period (Novelo et al., 2023). Prolonged viral detection up to 21 days post-infection, combined with sustained adult survival under temperate conditions, indicates that *Ae. koreicus* can remain infectious over extended periods and across multiple gonotrophic cycles.

Collectively, these findings demonstrate the potential for *Ae. koreicus* to support sustained arbovirus transmission in temperate environments, with transmission intensity shaped by temperature. Because vector capacity depends on the interaction between mosquito traits and environmental conditions, even modest physiological competence can translate into substantial transmission potential when ecological factors align. In particular, high vector abundance can amplify transmission potential even when intrinsic competence is moderate (Cator et al., 2020; Scott & Morrison, 2010). Within this framework, the combination of favourable thermal conditions and increasing population densities may enable *Ae. koreicus* to contribute to transmission, particularly in spatially localised outbreak scenarios. The convergence of prolonged adult survival, opportunistic anthropophilic behaviour, and the potential for high seasonal abundance suggests that *Ae. koreicus* could influence the timing and intensity of arbovirus transmission under moderate thermal conditions typical of temperate regions (Montarsi et al., 2022). This raises the possibility that, rather than driving large-scale epidemics, *Ae. koreicus* may act as a secondary or complementary vector, modulating transmission dynamics in expanding ecological niches under ongoing climate change, while also contributing to localized nuisance burdens.

## Acknowledgments

We thank Carolina Aguzzi for supporting with insectary work and mosquito dissections, and Giovanni Marini for fruitful discussions.

## Reference

ARPAV. (2023). Temperatura – anno 2023. Agenzia Regionale per la Prevenzione e Protezione Ambientale del Veneto. https://www.arpa.veneto.it/arpavinforma/indicatori-ambientali/indicatori_ambientali/clima-e-rischi-naturali/clima/temperatura/2023

Ballardini, M., Ferretti, S., Chiaranz, G. et al. First report of the invasive mosquito Aedes koreicus (Diptera: Culicidae) and of its establishment in Liguria, northwest Italy. Parasites Vectors 12, 334 (2019). 10.1186/s13071-019-3589-2

Blaha, M., Matiu, M., Majone, B., Zardi, D., Flacio, E., Anicic, N., Müller, G., Cherix, D., Gschwind, M., Modespacher, B., Müller, P., Erndle, K., Rizzoli, A., Stenico, A., Cassina, F., Rosà, R., & Da Re, D. (2026). Climate-change-driven shifts in the population dynamics of the invasive tiger mosquito (Aedes albopictus) in the Alps. 10.32942/X2T952

Briere, J. F., Pracros, P., Le Roux, A. Y., & Pierre, J. S. (1999). A Novel Rate Model of Temperature-Dependent Development for Arthropods. Environmental Entomology, 28(1), 22–29. 10.1093/ee/28.1.22

Brustolin, M., Bartholomeeusen, K., Rezende, T., Ariën, K., Müller, R. (2024) Mayaro virus, a potential threat for Europe: vector competence of autochthonous vector species. Parasites & Vectors 14:200 10.1186/s13071-024-06293-7

Brustolin M, Pujhari S, Terradas G, Werling K, Asad S, Metz HC, Henderson CA, Kim D, Rasgon JL. In Vitro and In Vivo Coinfection and Superinfection Dynamics of Mayaro and Zika Viruses in Mosquito and Vertebrate Backgrounds. J Virol. 2023 Jan 31;97(1):e0177822. 10.1128/jvi.01778-22

Brooks, M. E., Kristensen, K., van Benthem, K. J., Magnusson, A., Berg, C. W., Nielsen, A., Skaug, H. J., Mächler, M., Bolker, B. M. (2017). glmmTMB balances speed and flexibility among packages for zero-inflated count data. The R Journal, 9(2), 378–400.

Burnham, K. P., Anderson, D. R. (2002). Model Selection and Multimodel Inference: A Practical Information-Theoretic Approach (2nd ed.). Springer.

Capelli, G., Drago, A., Martini, S., Montarsi, F., Soppelsa, M., Delai, N., Ravagnan, S., Mazzon, L., Schaffner, F., Mathis, A., Di Luca, M., Romi, R., & Russo, F. (2011). First report in Italy of the exotic mosquito species Aedes (Finlaya) koreicus, a potential vector of arboviruses and filariae. Parasites and Vectors, 4(1). 10.1186/1756-3305-4-188

Carlassara, M., Khorramnejad, A., Oker, H., Bahrami, R., Lozada-Chávez, A. N., Mancini, M. V., Quaranta, S., Body, M. J. A., Lahondère, C., & Bonizzoni, M. (2024). Population-specific responses to developmental temperature in the arboviral vector Aedes albopictus: Implications for climate change. Global Change Biology, 30(3). 10.1111/gcb.17226

Carrington, L. B., Armijos, M. V., Lambrechts, L., Barker, C. M., & Scott, T. W. (2013). Effects of Fluctuating Daily Temperatures at Critical Thermal Extremes on Aedes aegypti Life-History Traits. PLOS ONE, 8(3), e58824. 10.1371/journal.pone.0058824

Cator, L. J., Johnson, L. R., Mordecai, E. A., El Moustaid, F., Smallwood, T. R. C., LaDeau, S. L., Johansson, M. A., Hudson, P. J., Boots, M., Thomas, M. B., Power, A. G., & Pawar, S. (2020). The Role of Vector Trait Variation in Vector-Borne Disease Dynamics. Frontiers in Ecology and Evolution, 8. 10.3389/fevo.2020.0018

Ciocchetta, S., Darbro, J. M., Frentiu, F. D., Montarsi, F., Capelli, G., Aaskov, J. G., & Devine, G. J. (2017). Laboratory colonization of the European invasive mosquito Aedes (Finlaya) koreicus. Parasites and Vectors, 10(1). 10.1186/s13071-017-2010-2

Ciocchetta, S., Prow, N. A., Darbro, J. M., Frentiu, F. D., Savino, S., Montarsi, F., Capelli, G., Aaskov, J. G., & Devine, G. J. (2018). The new European invader Aedes (Finlaya) koreicus: a potential vector of chikungunya virus. Pathogens and Global Health, 112(3), 107–114. 10.1080/20477724.2018.1464780

Clark, J. S. (2003). Uncertainty and variability in demography and population growth: a hierarchical approach. Special Feature Ecology, 84(6), 1370–1381. 10.1890/0012-9658(2003)084[1370:UAVIDA]2.0.CO;2

Couper, L. I., Nalukwago, D. U., Lyberger, K. P., Farner, J. E., & Mordecai, E. A. (2024). How much warming can mosquito vectors tolerate? Global Change Biology, 30(12), 17610. 10.1111/gcb.17610

Craig MH, Snow RW, le Sueur D. A climate-based distribution model of malaria transmission in sub-Saharan Africa. Parasitol Today. 1999 Mar;15(3):105–11.https://www.cell.com/partod/abstract/S0169-4758(99)01396-4

Crowl, T. A., Crist, T. O., Parmenter, R. R., Belovsky, G., & Lugo, A. E. (2008). The spread of invasive species and infectious disease as drivers of ecosystem change. Frontiers in Ecology and the Environment, 6(5), 238–246. 10.1890/070151;PAGEGROUP:STRING:PUBLICATION

Da Re, D., Arnoldi, D., Rizzoli, A., & Marini, G. (2026). Cold tolerance and egg diapause shape overwintering success in Aedes koreicus mosquitoes. 10.2139/SSRN.6101679

Dennington, N. L., Grossman, M. K., Ware-Gilmore, F., Teeple, J. L., Johnson, L. R., Shocket, M. S., McGraw, E. A., & Thomas, M. B. (2024). Phenotypic adaptation to temperature in the mosquito vector, Aedes aegypti. Global Change Biology, 30(1), e17041. 10.1111/gcb.17041

Deutsch, C. A., Tewksbury, J. J., Huey, R. B., Sheldon, K. S., Ghalambor, C. K., Haak, D. C., & Martin, P. R. (2008). Impacts of climate warming on terrestrial ectotherms across latitude. Proceedings of the National Academy of Sciences of the United States of America, 105(18), 6668–6672. 10.1073/PNAS.0709472105;PAGE:STRING:ARTICLE/CHAPTER

Early, R., Bradley, B. A., Dukes, J. S., Lawler, J. J., Olden, J. D., Blumenthal, D. M., Gonzalez, P., Grosholz, E. D., Ibañez, I., Miller, L. P., Sorte, C. J. B., & Tatem, A. J. (2016). Global threats from invasive alien species in the twenty-first century and national response capacities. Nature Communications 2016 7:1, 7(1), 12485-. 10.1038/ncomms12485

Lafferty, K. D. (2009). The ecology of climate change and infectious diseases. Ecology,90(4), 888– 900. 10.1890/08-0079.1

Folguera, G., Mensch, J., Munoz, JC eballos, S.G., Hasson, E., Bozinovic, F (2010). Ontogenetic stage-dependent effect of temperature on developmental and metabolic rates in a holometabolous insect. Journal of Insect Physiology, 56(11).10.1016/j.jinsphys.2010.06.015

Ghasemi A, Zahediasl S. Normality tests for statistical analysis: a guide for non-statisticians. Int J Endocrinol Metab. 2012 Spring;10(2):486–9. doi: 10.5812/ijem.3505. Epub 2012 Apr 20. 10.5812/ijem.3505

Gulia-Nuss M, Eum JH, Strand MR, Brown MR. Ovary ecdysteroidogenic hormone activates egg maturation in the mosquito Georgecraigius atropalpus after adult eclosion or a blood meal. J Exp Biol. 2012 Nov 1;215(Pt 21):3758–67. 10.1242/jeb.074617

Hartig, F. (2016). DHARMa: Residual Diagnostics for Hierarchical (Multi-Level / Mixed) Regression Models. CRAN: Contributed Packages. 10.32614/CRAN.package.DHARMa

Heitmann A, Jansen S, Lühken R, Leggewie M, Schmidt-Chanasit J, Tannich E. Forced salivation as a method to analyze vector competence of M=mosquitoes. J Vis Exp. 2018 Aug 7;(138):57980. 10.3791/57980

Huey, R. B., Deutsch, C. A., Tewksbury, J. J., Vitt, L. J., Hertz, P. E., Pérez, H.J.Á., & Garland, T. (2009). Why tropical forest lizards are vulnerable to climate warming. Proceedings of the Royal Society B: Biological Sciences, 276(1664), 1939–1948. 10.1098/RSPB.2008.1957

Jansen, S., Cadar, D., Lühken, R., Pfitzner, W. P., Jöst, H., Oerther, S., Helms, M., Zibrat, B., Kliemke, K., Becker, N., Vapalahti, O., Rossini, G., Heitmann, A. (2021). Vector Competence of the Invasive Mosquito Species Aedes koreicus for Arboviruses and Interference with a Novel Insect Specific Virus. Viruses 2021, Vol. 13, 13(12). 10.3390/V13122507

Jarosík V, Kratochvíl L, Honek A, Dixon AF. A general rule for the dependence of developmental rate on temperature in ectothermic animals. Proc Biol Sci. 2004 May 7;271 Suppl 4(Suppl 4):S219–2. 10.1098/rsbl.2003.0145

Kandethody Ramachandran, Chris P Tsokos, B.M., York, N., & Diego, S. (2020). Mathematical statistics with applications in R. https://books.google.com/books?hl=en&lr=&id=t3bLDwAAQBAJ&oi=fnd&pg=PP1&ots=1tlPNWeRRG&sig=vHI7PgUqosW9IDYMAfUBzRuOhH4

enneth L. Knight, The Aedes (Finlaya) Chrysolineatus Group of Mosquitoes (Diptera: Culicidae)(1947), Annals of the Entomological Society of America, 40(4), 624–649, 10.1093/aesa/40.4.624

Kishore, J., Goel, M., & Khanna, P. (2010). Understanding survival analysis: Kaplan-Meier estimate. International Journal of Ayurveda Research, 1(4), 274–278.

Lee, S., & Lee, D. K. (2018). What is the proper way to apply the multiple comparison test? Korean Journal of Anesthesiology, 71(5). 10.4097/kja.d.18.00242

Marcantonio, M., Reyes, T., & Barker, C. M. (2019). Quantifying Aedes aegypti dispersal in space and time: a modelling approach. (2019).Wiley Online Library, 10(12). 10.1002/ecs2.2977

Marini G, Arnoldi D, Baldacchino F, Capelli G, Guzzetta G, Merler S, Montarsi F, Rizzoli A, Rosà R. First report of the influence of temperature on the bionomics and population dynamics of Aedes koreicus, a new invasive alien species in Europe. Parasit Vectors. 2019 Nov 6;12(1):524. 10.1186/s13071-019-3772-5

Martens, W. J. M., Jetten, T. H., & Focks, D. A. (1997). Sensitivity of malaria, schistosomiasis and dengue to global warming. SpringerWJM Martens, TH Jetten, DA FocksClimatic Change, 1997•Springer, 35(2), 145–156. 10.1023/A:1005365413932

Miyagi, I. (1971). Notes on the Aedes (Finlaya) chrysolineatus subgroup in Japan and Korea (Diptera: Culicidae). Tropical Medicine, 13(3). https://nagasaki-u.repo.nii.ac.jp/records/24725

Montarsi, F., Martini, S., Dal Pont, M., Delai, N., Ferro Milone, N., Mazzucato, M., Soppelsa, F., Cazzola, L., Cazzin, S., Ravagnan, S., Ciocchetta, S., Russo, F., & Capelli, G. (2013). Distribution and habitat characterization of the recently introduced invasive mosquito Aedes koreicus [Hulecoeteomyia koreica], a new potential vector and pest in north-eastern Italy. Parasites & Vectors, 6(1), 292. 10.1186/1756-3305-6-292

Montarsi F, Rosso F, Arnoldi D, Ravagnan S, Marini G, Delucchi L, Rosà R, Rizzoli A. First report of the blood-feeding pattern in Aedes koreicus, a new invasive species in Europe. Sci Rep. 2022 Sep 21;12(1):15751. 10.1038/s41598-022-19734-z

Mordecai, E.A., Caldwell, J.M., Grossman, M.K., Lippi, C.A., Johnson, L.R., Neira, M., Rohr, J.R., Ryan, S.J., Savage, V., Shocket, M.S., Sippy, R., Stewart Ibarra, A.M., Thomas, M.B., and Villena, O. (2019). Thermal biology of mosquito-borne disease. Ecol Lett, 22: 1690–1708. 10.1111/ele.13335

Novelo, M., Dutra, H. L. C., Metz, H. C., Jones, M. J., Sigle, L. T., Frentiu, F. D., Allen, S. L., Chenoweth, S. F., & McGraw, E. A. (2023). Dengue and chikungunya virus loads in the mosquito Aedes aegypti are determined by distinct genetic architectures. PLOS Pathogens, 19(4), e1011307. 10.1371/journal.ppat.1011307

Overgaard, J., Kearney, M. R., & Hoffmann, A. A. (2014). Sensitivity to thermal extremes in Australian Drosophila implies similar impacts of climate change on the distribution of widespread and tropical species. Global Change Biology, 20(6), 1738–1750. 10.1111/GCB.12521;JOURNAL:JOURNAL:13652486;WGROUP:STRING:PUBLICATION

Padfield, D., & Matheson, G. (2020). nls.multstart: Robust non-linear regression using AIC scores. R package version 1.2.0. https://cran.r-project.org/package=nls.multstart

Padfield, D., O Sullivan, H., & amp; Pawar, S. (2021). rTPC and nls.multstart: A new pipeline to fit thermal performance curves in R. Methods in Ecology and Evolution, 12(6), 1138–1143.

Parham, P.E. and Michael, E. (2010) Modeling the Effects of Weather and Climate Change on Malaria Transmission. Environmental Health Perspectives, 118, 620–626.10.1289/ehp.0901256

Parmesan, C. (2006). Ecological and evolutionary responses to recent climate change. Annual Review of Ecology, Evolution, and Systematics, 37(Volume 37, 2006), 637–669. 10.1146/ANNUREV.ECOLSYS.37.091305.110100/CITE/REFWORKS

Pawar, S., Huxley, P. J., Smallwood, T. R. C., Nesbit, M. L., Chan, A. H. H., Shocket, M. S., Johnson, L. R., Kontopoulos, D. G., & Cator, L. J. (2024). Variation in temperature of peak trait performance constrains adaptation of arthropod populations to climatic warming. Nature Ecology and Evolution, 8(3), 500–510. 10.1038/s41559-023-02301-8

Pawar, S., & Kontopoulos, D. G. (2025). Toward a general understanding of thermal performance curves in biology. Proceedings of the National Academy of Sciences of the United States of America, 122(51), e2528528122. 10.1073/pnas.2528528122

Ramachandran, K. M., & Tsokos, C. P. (2020). Mathematical statistics with applications in R (3rd ed.). Academic Press.

R Core Team. (2022). R Core Team 2021 R: A language and environment for statistical computing. R foundation for statistical computing. https://www.R-project.org/. R Foundation for Statistical Computing, 2.

Rabitsch, W., Essl, F., & Schindler, S. (2017). The Rise of Non-native Vectors and Reservoirs of Human Diseases. Impact of Biological Invasions on Ecosystem Services, 263–275. 10.1007/978-3-319-45121-3_17

Rebolledo AP, Sgrò CM and Monro K (2021) Thermal Performance Curves Are Shaped by Prior Thermal Environment in Early Life. Front. Physiol. 12:738338. 10.3389/fphys.2021.738338

Reuss, F., Wieser, A., Niamir, A., Bálint, M., Kuch, U., Pfenninger, M., Müller, R. (2018) Thermal experiments with the Asian bush mosquito (Aedes japonicus japonicus) (Diptera: Culicidae) and implications for its distribution in Germany. Parasites & Vectors 11:81

Robert, L. L., & Olson, J. K. (1998). Effects of sublethal dosages of insecticides on Culex quinquefasciatus. Journal of the American Mosquito Control Association, 5, 239–246.

Schneider, J., Valentini, A., Dejean, T., Montarsi, F., Taberlet, P., Glaizot, O., & Fumagalli, L. (2016). Detection of Invasive Mosquito Vectors Using Environmental DNA (eDNA) from Water Samples. PLOS ONE, 11(9), e0162493. 10.1371/journal.pone.0162493

Scott, T. W., & Morrison, A. C. (2010). Vector dynamics and transmission of dengue virus: implications for dengue surveillance and prevention strategies: vector dynamics and dengue prevention. Current Topics in Microbiology and Immunology, 338, 115–128. 10.1007/978-3-642-02215-9_9

Sinclair BJ, Marshall KE, Sewell MA, Levesque DL, Willett CS, Slotsbo S, Dong Y, Harley CD, Marshall DJ, Helmuth BS, Huey RB. Can we predict ectotherm responses to climate change using thermal performance curves and body temperatures? Ecol Lett. 2016 Nov;19(11):1372–1385. 10.1111/ele.12686

Slayi, M., & Jaja, I. F. (2025). Strategies for mitigating heat stress and their effects on behavior, physiological indicators, and growth performance in communally managed feedlot cattle. Frontiers in Veterinary Science, 12, 1513368. 10.3389/fvets.2025.1513368

Vanslembrouck, A., Jansen, S., De Witte, J., Janssens, C., Vereecken, S., Helms, M., Lange, U., Lühken, R., Schmidt-Chanasit, J., Heitmann, A., Müller, R. (2024) Larval competition between Aedes and Culex mosquitoes carries over to higher arboviral infection during their adult stage. Viruses 16:1202

Versteirt, V., De Clercq, E. M., Fonseca, D. M., Pecor, J., Schaffner, F., Coosemans, M., Van Bortel, W. (2012). Bionomics of the Established Exotic Mosquito Species Aedes koreicus in Belgium, Europe, Journal of Medical Entomology, 49(6),1226–1232, 10.1603/ME11170

WeatherSpark. (2026). Average spring weather in Europe. Retrieved April 3, 2026, from https://weatherspark.com/

World Health Organization (2025) Dengue and severe dengue. Available at: https://www.who.int/news-room/fact-sheets/detail/dengue-and-severe-dengue (accessed 31 March 2026).

Williges, E., Farajollahi, A., Scott, J. J., Mccuiston, L. J., Crans, W. J., & Gaugler, R. (2008). Scientific note laboratory colonization of Aedes japonicus. J Am Mosq Control Assoc. 2008 Dec;24(4):591–3. 10.2987/5714.1

Yin, X., Kropff, M., McLaren, G., Forest, R. V.-A. and, & 1995, undefined. (1995). A nonlinear model for crop development as a function of temperature. Elsevier, 77, 1–6. https://www.sciencedirect.com/science/article/pii/016819239502236Q

