## Supplementary Material for "Thermal responses of an emerging temperate mosquito reshape arboviral transmission risk"

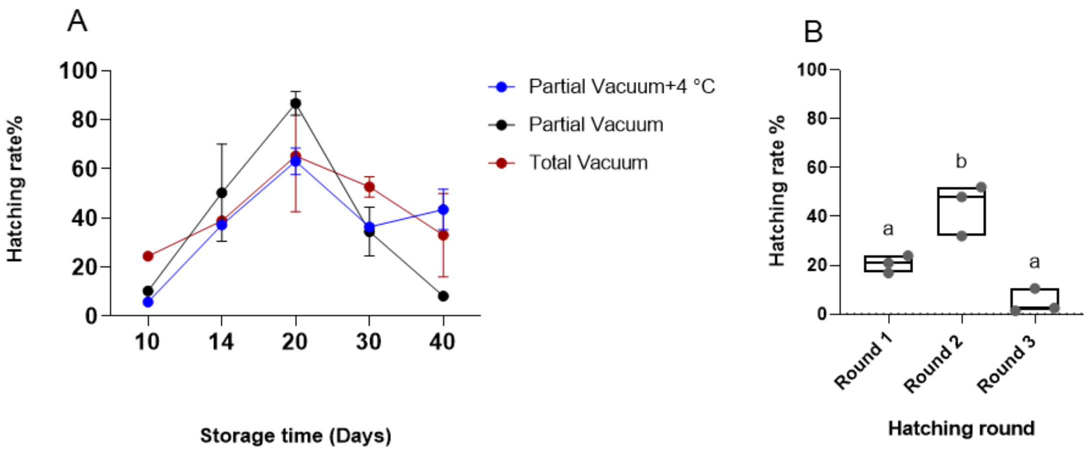


Figure S1. **Hatching performance of *Ae. koreicus* eggs.** (A) Hatching rate of *Ae. koreicus* eggs after different quiescent periods. At each time point, the median value from 2 biological replicates was shown. Error bars represent ±SE. (B) Hatching rate of *Ae. koreicus* eggs at different rounds of hatching. (B) Hatching rate (%) across three consecutive hatching rounds (Round 1–3). Different lowercase letters indicate significant differences among rounds ($p < 0.05$). Boxplots show median and interquartile range; points represent biological replicates.

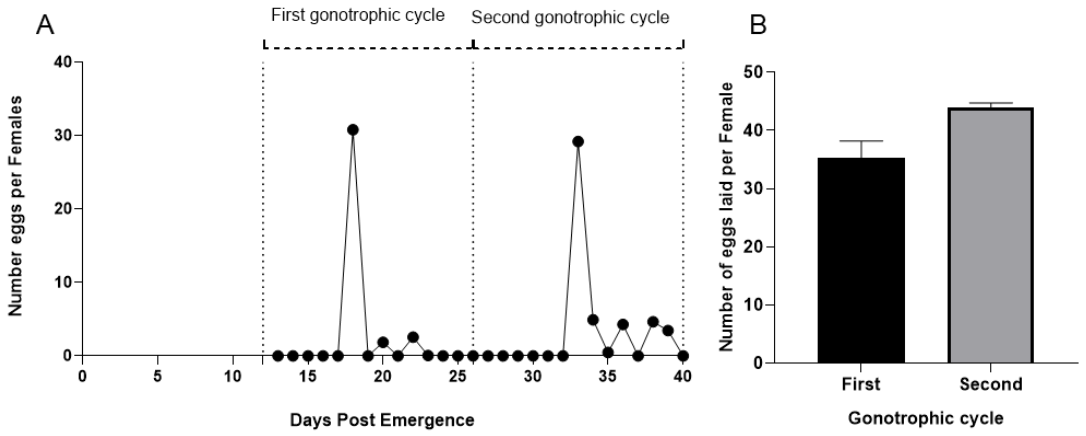


Figure S2 **Reproductive output of *Ae. koreicus* females across two gonotrophic cycles.** (A) Gonotrophic cycles showing egg production per blood-feeding event. Each dot represents the mean daily number of eggs laid from two biological replicates; horizontal dashed lines indicate blood-feeding events. (B) Comparison of reproductive output between the first and second gonotrophic cycles, expressed as the mean total number of eggs laid over the entire cycle from two biological replicates.

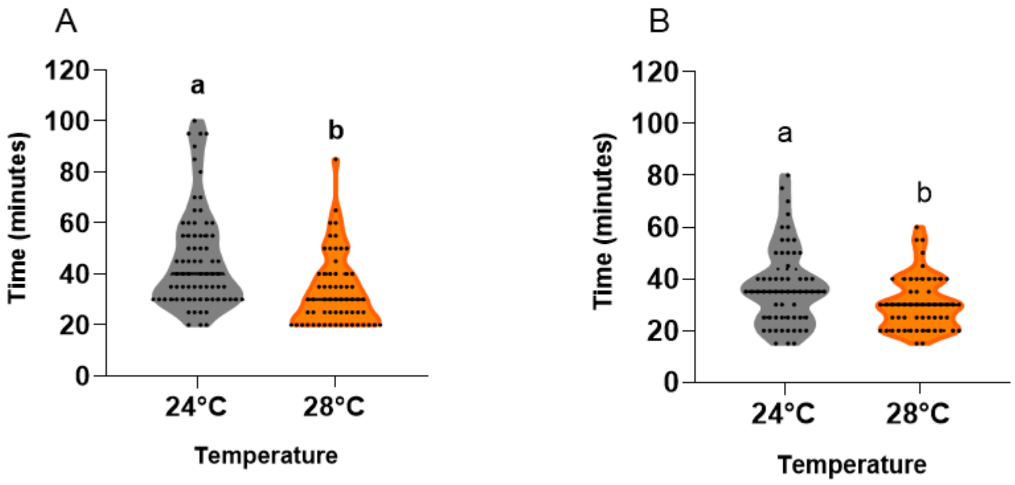


Figure S3. **Knockdown time of *Ae. koreicus* under cold thermal stress (4 °C) following rearing at different temperatures.** (A) Knockdown time of males. (B) Knockdown time of females. Each dot represents an individual mosquito. Different lowercase letters indicate significant differences among treatments ($p < 0.05$, Student's t-test).

Table S1. Sample sizes used for vector competence assays

| Time points (DPI) | 24 °C–CHIKV(n) | 28 °C–CHIKV(n) | 24 °C–DENV(n) | 28 °C–DENV(n) |
|---|---|---|---|---|
| 7 DPI | 18 | 20 | 26 | 17 |
| 14 DPI | 17 | 19 | 27 | 20 |
| 21DPI | 25 | 36 | 26 | 26 |

Number of mosquitoes analysed for each combination of temperature, virus, and time point (dpi) in vector competence assays.

Table S2. Thermal performance parameters for three life-history traits of *Ae. koreicus*

| Traits | Model | $CT^{TIn}$ (°C) | $T_{oht}$ (°C) | $CT^{TIx}$ (°C) | Notes |
|---|---|---|---|---|---|
| Developmental success (Larvae to adult survival) | Beta-binomial (glmmTMB) | _ | 26.8 | _ | Thermal limit not estimated |
| Developmental speed | Brière-2 (NLS) | 13.2 | 29.6 | 35.0* | $CT_{min}$: model |
| Adult longevity (Females) | Weibull AFT (flexsurv) | _ | 20.6 | _ | Censored individuals are included |
| Adult longevity (Males) | Weibull AFT (flexsurv) | _ | 20.8 | _ | Censored individuals are included |

CT min =lower critical thermal limit; T opt = thermal optimum; CT max = upper critical thermal limit. —indicates parameter not estimable from the fitted model. *CT max for development speed is the observed experimental upper bound (no development at 35°C) rather than a freely estimated model parameter.
